# Memory entrainment by visually evoked theta-gamma coupling

**DOI:** 10.1101/191189

**Authors:** Moritz Köster, Ulla Martens, Thomas Gruber

**Author notes:** Corresponding author: Moritz Köster, Free University Berlin, Institute of Psychology, Habelschwerdter Allee 45, 14195 Berlin, Germany.

## Abstract

It is an integral function of the human brain to sample novel information from the environment and to update the internal representation of the external world. The formation of new memories is assumed to be orchestrated by neuronal oscillations, the rhythmic synchronization of neuronal activity within and across cell assemblies. Specifically, successful encoding of novel information is associated with increased theta oscillations (3-8Hz) and theta coupled gamma activity (40-120Hz), and a decrease in alpha oscillations (8-12Hz). However, given the correlative nature of neurophysiological recordings, the causal role of neuronal rhythms in human memory encoding is still unclear. Here, we experimentally enhance the formation of novel memories by a visual brain stimulation at an individually adjusted theta frequency, in contrast to the stimulation at an individual alpha frequency. Critically, the memory entrainment effect by the theta stimulation was not explained by theta power *per se*, but was driven by visually evoked theta-gamma coupling in wide spread cortical networks. These findings provide first evidence for a functional role of the theta rhythm and the theta-gamma neuronal code in human episodic memory. Yet more strikingly, the entrainment of mnemonic network mechanisms by a simplistic visual stimulation technique provides a proof of concept that internal rhythms align with visual pacemakers, which can entrain complex cognitive functions in the wake human brain.

## Introduction

To retain a coherent internal representation of the outer world, the wake human brain constantly samples and integrates novel information from the environment (1, 2), a process that accompanies perception (3). How is this complex function accomplished and which are the neuronal mechanisms underlying memory encoding? The rhythmic synchronization of neuronal activity within and across nerve cell populations is posited to coordinate and integrate distributed processes across the brain (4–6) and assumed to be a key mechanism underpinning the real-time integration of novel experiences into existing neuronal representations (7, 8).

In subsequent memory paradigms, the neuronal activity during encoding is contrasted between later remembered and later forgotten items to isolate the neuronal processes underlying memory formation (9). Using electro- and magnetoencephalography, it has been established that the formation of novel memory traces is closely associated with neuronal oscillatory activity in the theta (3-8 Hz), alpha (8-12 Hz) and gamma (40-120 Hz) frequency in the human brain (10–12). Specifically, successful encoding of visual stimuli was marked by increases in theta and gamma power and a decrease in alpha power (11, 12).

Theta oscillations are proposed to serve the ordering and binding of perceptual information, which are reflected in gamma oscillations (13, 14), forming a theta-gamma neuronal code (15). This is substantiated by theta-gamma phase-amplitude coupling (PAC) pattern in the human neocortex (16), and an increase in theta-gamma PAC in cortical and medio-temporal networks during successful memory encoding (12, 17, 18). The theta-gamma code is assumed to facilitate the integration of perceptual information into existing networks by squeezing real time events onto a neuronal time scale (19) to support long-term potentiation processes in the hippocampus (20), the core system of human episodic memory (21, 22). Reduced alpha oscillations during successful encoding may reflect attention gating processes during encoding (23, 24).

Critically, empirical findings that fuel contemporary theoretical models on distinct roles of neuronal theta, alpha and gamma oscillations in memory encoding are foremost correlational. This is, oscillatory activity observed in the human brain could be an epiphenomenal byproduct of perceptual and mnemonic processes in neuronal networks, without a causal function (19). There is first evidence that memory consolidation processes during sleep can be experimentally enhanced by transcranial alternating currents (25) and auditory stimulation (26) in a slow-wave delta rhythm (0.5 – 2 Hz). Furthermore, transcranial alternating current stimulation at the theta frequency were successfully applied to enhance working memory (27). Yet, the causal relevance of oscillatory dynamics during memory encoding in the wake human brain is still unclear.

In the present study, we experimentally manipulated the memory encoding processes by a visual stimulation at an individually adjusted theta rhythm to selectively enhance the memory encoding process, compared to a stimulation at an individual alpha frequency, which may disrupt encoding. A scalp electroencephalogram (EEG) was recorded to measure the entrainment of theta, alpha and phase-amplitude coupling (PAC) dynamics and to test the impact of the visual entrainment on subsequent memory performance.

## Results

### Visual theta and alpha stimulation effect subsequent memory performance

Subjects (n = 20) saw visually flickering stimuli, presented at an individual theta or alpha frequency, using the duty-cycles (i.e., on and off screens) of an CTR monitor (Figure 1A). Importantly, a pre-study was conducted to adjust theta and alpha frequencies individually (12), due to strong inter-individual variability in the theta and the alpha frequency (10). We further presented static stimuli to estimate the difference in memory performance for flickering and non-flickering stimuli. In a retrieval session, we assessed how the visual stimulation during encoding effected subjects subsequent memory performance (SME = percent of subsequently remembered [SR] minus subsequently forgotten [SF] items).

The stimulation at an individual theta frequency increased subsequent memory performance (SME), compared to the alpha stimulation, *t*(19) = 2.24, *p* = .037, Cohen’s *d* = .50 (see Figure 1B), hereafter called memory entrainment effect. This was due to a higher number of subsequently remembered (SR) items in the theta stimulation condition, compared to the alpha stimulation, 23.8 % vs. 21.5 %, *t*(19) = 2.13, *p* = .046, as well as a higher rate of subsequently forgotten (SF) items in the alpha stimulation condition, compared to the theta stimulation, 47.4 % vs. 49.9 % *t*[19] = 1.99, *p* = .062. As expected, static pictures were remembered best (SR: 28.2 % and SF: 42.6 %), with a higher SME compared to the theta (*t*[19] = 4.15, *p* < .001), and the alpha stimulation (*t*[19] = 10.97, *p* < .001). The stimulation condition did not affect the rates of subsequent know (SK) responses (theta: 28.7 %, alpha: 28.6 %, static: 29.3 %). Note that the mean differences for SR and SF were calculated post hoc, following significant main effects and interactions in the overall ANOVA (3 Conditions [theta, alpha, gamma] x 3 Responses [SR, SK, SF]) and the subsidiary ANOVAs for SR and SF responses, all *ps* < .001.

**Figure 1.**
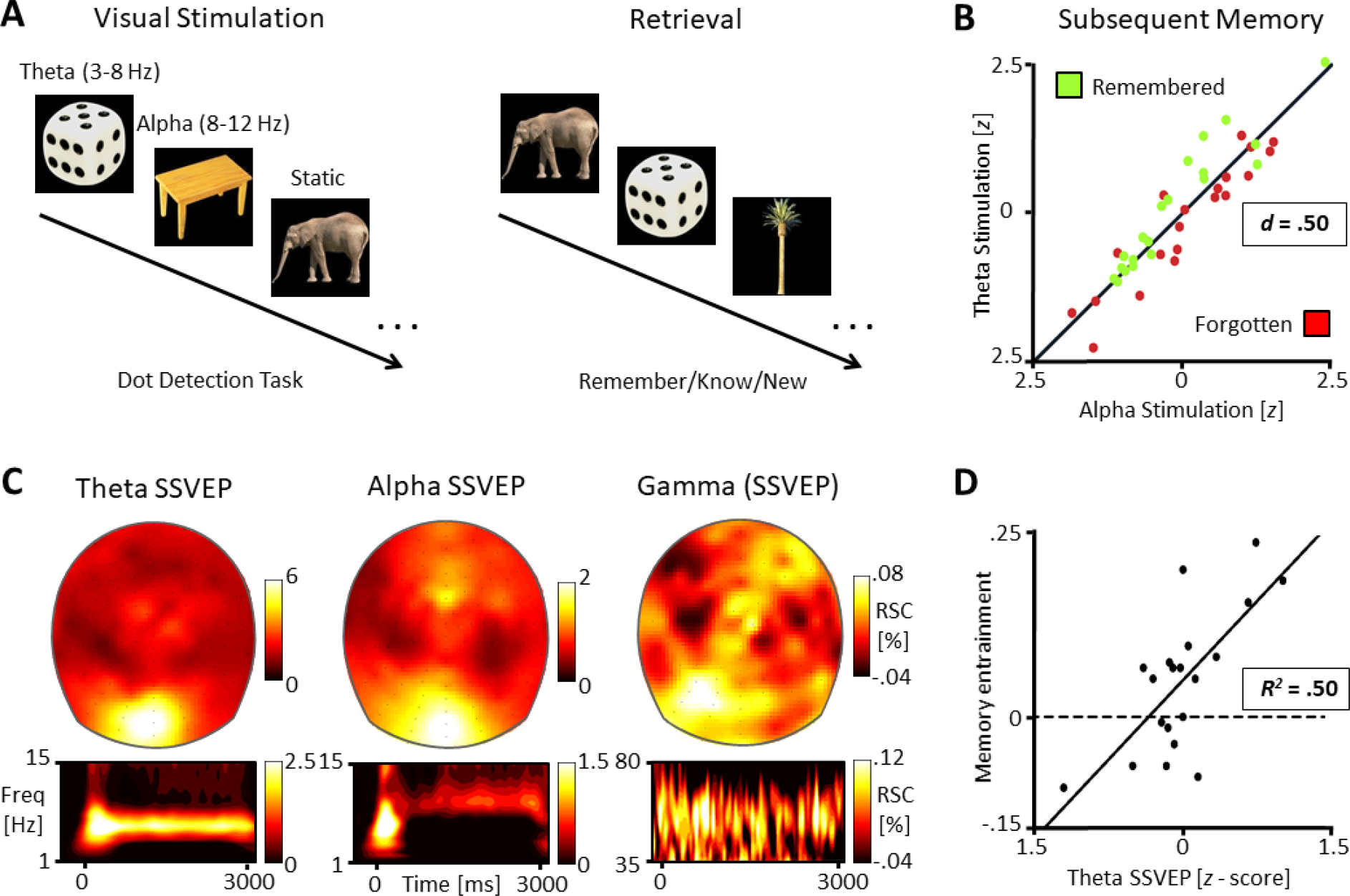
Study design and visual stimulation effects on subsequent memory performance. (A) Object pictures were presented at different stimulation frequencies; individually adjusted theta (3-8 Hz), alpha (8-12 Hz), or static (non-flickering). (B) Subsequently remembered (SR) and forgotten (SF) response rates are displayed (z-standardized for the combined visualization). Each subject is represented by a green and a red dot; Green dots above the diagonal indicate subjects with a higher SR rate for theta, red dots below the diagonal indicate subjects with a higher SF rate for alpha. The subsequent memory performance (SME = SR - SF) was higher for theta, compared to alpha (*p* < .05, Cohen’s *d* = .50). (C) Topographies display the steady state visually evoked potentials (SSVEP) at the theta and alpha frequencies. The gamma topography displays the 50–80 Hz activity accompanying theta and alpha SSVEPs. Values indicate relative signal changes (RSC, 500-3000ms). Time-frequency plots display the spectral power at the topographical peak electrode. (D) Inter-individual differences in memory entrainment (Theta SME – Alpha AME) were explained by the participants theta SSVEP (*r* = .71, *p* < .001, *R^2^* = .50), but not by alpha SSVEP (*p* = .697; not displayed).

The effect size of the memory entrainment effect was at a reasonable, intermediate level. However, not all participants were equally responsive to the visual stimulation (i.e., participants close to the 45° diagonal in Figure 1B). Thus, in a next step, we tested if interindividual differences in participants brain response to the visual stimulation would explain differences in the memory entrainment effect (theta SME – alpha SME).

### Participants responsiveness to the theta stimulation predicts memory entrainment

Visually flickering stimuli elicit steady state visually evoked potentials (SSVEPs), in the scalp recorded EEG (28). A SSVEP is the oscillatory response of the visual cortex to a rapidly repeating (flickering) stimulus, at the specific frequency of the driving stimulus. Here, the neuronal responses to the visual theta and alpha stimulation were recorded at 128 EEG electrodes.

The theta and the alpha stimulation elicited clear SSVEP signals over the visual cortex, accompanied by increased gamma oscillations at posterior recording sites, all *t*(19) > 3.11, *p* < .006 (Figure 1C). In a first step, we tested if inter-individual differences in memory entrainment were due to differences in participants responsiveness to the visual stimulation. The responsiveness to the visual stimulation was quantified by the power at the peak electrode and the spread of the SSVEP signal, determined by the entropy. Memory entrainment was clearly predicted by the power and spread of the theta SSVEP, *r* = .71, *p* = .00043, *R^2^* = .50 (see Figure 1D), but no relation was found between the alpha SSVEP signal and the memory entrainment effect, *r* = .09, *p* = .697. Thus, memory entrainment was driven by memory enhancing effects of the theta stimulation, but not by disrupting effects of the alpha stimulation.

### Memory entrainment is driven by visually evoked theta-gamma phase-amplitude coupling (PAC)

In a second step, we tested the neuronal dynamics during encoding that predict subsequent memory performance, by comparing the neuronal activity between encoding conditions. Surprisingly, although theta SSVEPs explained inter-individual differences in memory entrainment, theta oscillations showed an inverse SME, namely higher theta SSVEPs for SF, than SK, than SR items, at the peak electrode (Figure 2A, upper panel), *F*(2, 38) = 6.41, *p* = .004. This inverse SME was distributed across posterior, central and frontal regions (Figure 2A, lower panel). The memory enhancing effect of the theta stimulation was resolved by the theta-gamma PAC analyses, testing the coupling between phase of the individual theta SSVEP, at the theta peak electrode, and the 50-80Hz gamma amplitude (Figure 2B), at all 128 electrodes, quantified by the modulation index (MI) (16). The averaged MI cross all 128 electrodes, revealed a clear SME, namely higher theta-gamma PAC for SR, compared to SK, compared to SF items, *F*(2, 38) = 8.08, *p* < .001. The difference in theta-gamma PAC between SR and SF stimuli was spread in wide cortical networks and was highest at left temporal and centro-frontal electrodes (Figure 2B).

**Figure 2.**
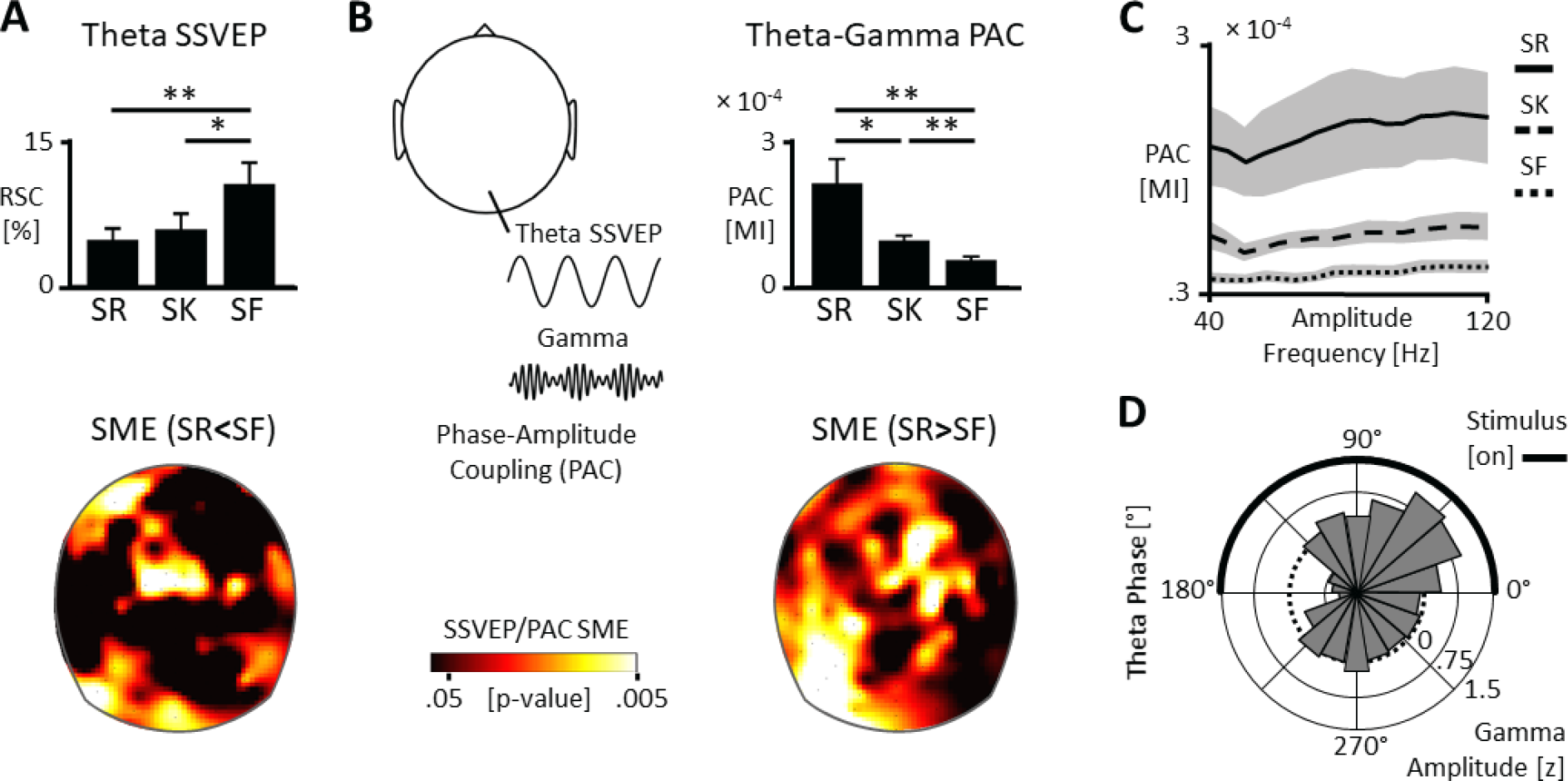
Memory entrainment by theta-gamma coupling. (A) Bars indicate the theta SSVEP signal differences between subsequently remembered (SR), known (SK), and forgotten (SF) items, at the peak electrode (ANOVA: *F*[2, 38] = 6.41, *p* = .004). The topographical map illustrates the distribution of the inverse subsequent memory effect (SME, SF > SR) across the scalp. Note that alpha SSVEP and gamma did not differ between conditions at the peak electrode. Significance levels: \**p* < .05, \*\**p* < .01. (B) The cartoon head illustrates theta-gamma phase amplitude coupling, as assessed by the modulation index (MI). Bars display the mean differences in MI, averaged across all 128 electrodes (*F*[2, 38] = 8.08, *p* < .001). The topographical map displays the distribution of the SME (SR > SF) in theta-gamma PAC across the 128 scalp electrodes. Note that overall theta-gamma PAC was much higher than overall alpha-gamma PAC (*p* < .002, not displayed). (C) Theta-gamma PAC across the 40-120Hz gamma range. MI values indicate the mean PAC between gamma amplitude at each of 128 electrodes to the individual SSVEP theta phase at the peak electrode. (D) The circular histogram displays subsequent memory differences (SR-SF) for z-standardized gamma amplitudes at the left temporal peak electrodes of the MI difference plot.

Further inspection of theta-gamma PAC revealed that the SME in theta-gamma coupling pattern was not restricted to the preselected gamma frequency range (50-80Hz), but was present across a broad gamma range, from 40 to 120Hz (Figure 2C). We further looked at the critical theta phase of theta-gamma PAC for successful memory encoding at left temporal recording sites. The gamma amplitude difference (SR-SF), was specifically high at an early theta phase (0°-90°), corresponding to the onset of the flickering stimulus, but lower towards the offset of the flickering stimulus (120°-210°; Figure 2D).

No differences between encoding conditions (SR, SK, SF) were found for the alpha SSVEPs, *F*(2, 38) = 0.74, *p* = .486, and the gamma power accompanying the SSVEP conditions (displayed in Figure 1C), when tested in an ANOVA with all subsequent responses (SR, SK, SF) and both SSVEP conditions (theta and alpha), all *ps* > .634, for all main effects and interactions. In addition, across all conditions and all electrodes, theta-gamma PAC was much higher than alpha-gamma PAC, *t*(19) = 3.53, *p* = .002. Thus, neither the alpha SSVEPs nor the accompanying gamma power as such differed between encoding conditions.

To conclude, theta SSVEPs were higher for SF compared to SR stimuli, whereas theta-gamma PAC was higher for SR compared to SF stimuli. Thus, the memory entrainment effect of theta stimulation was not explained by theta SSVEP power *per se*, but by theta-gamma PAC processes. Note that the inverse SME for theta power (SR < SF) rules out the possibility that the positive SME in theta-gamma PAC (SR > SF) could be due to differences in theta SSVEPs power between conditions.

## Discussion

The present results provide first evidence for a functional role of the theta rhythm and the theta-gamma neuronal code in human episodic memory (15). Specifically, subsequent memory performance was higher for a visual brain stimulation at an individual theta frequency, compared to a stimulation at an individual alpha frequency. This memory entrainment effect was strongest for participants with a high responsiveness to the theta stimulation, indicating that memory entrainment was due to entrainment effects of the theta stimulation. However, subsequent memory performance was not explained by the theta SSVEP *per se*, but was driven by theta-gamma PAC in distributed cortical networks. These results provide a proof of concept that simplistic external pacemakers can stimulate rhythmic processes in the wake human brain and thereby entrain complex cognitive functions, such as memory encoding.

The alpha SSVEP stimulation was associated with a lower memory performance, than the theta stimulation. However, the alpha SSVEP signal did not explain inter-individual differences in memory entrainment and did not dissociate between encoding conditions (SR, SK, SF). Furthermore, the alpha SSVEP elicited much lower alpha-gamma PAC pattern, when compared to theta-gamma PAC. Thus, the effect of alpha SSVEPs was unspecific compared to the visually entrained theta rhythm and theta-gamma PAC pattern. As expected, non-flickering stimuli were remembered best. This may be due to the physically longer presentation time (no off frames), or because, in the absence of external stimulation, the brain could more flexibly adjust its rhythmicity.

Intuitively, one could interpret the present results in terms of a slower (3-8Hz) versus a faster (8-12Hz) pace of stimulus presentation. However, since the absolute presentation times were matched between the two stimulation conditions (theta and alpha), this would lead to the same conclusion, namely that a presentation at the theta pace is more favorable for mnemonic processing than a presentation at the alpha pace. Importantly, besides the memory entrainment at a behavioral level, recording a high-density EEG, we could identify the neuronal mechanism that explained differences in memory entrainment between subjects and at the subject level. This is, the theta SSVEP strength explained differences in memory entrainment between subjects, and theta-gamma PAC was higher for subsequently remembered, compared to subsequently forgotten items.

Theta oscillations in the cerebral cortex are associated with mnemonic control processes, namely the ordering and binding of perceptual information, reflected in gamma oscillations, which are nested in the theta phase (13–15, 17). In line with this idea, we found a strong subsequent memory effect in theta-gamma PAC and the theta stimulation selectively improved context-dependent, associative memories (SR, “remember” responses), but not familiarity based memory processes (SK, “know” responses) (29). However, while the theta rhythm is ascribed a prefrontal and medio-temporal control function in human memory processes (2, 30), the present study adds to recent evidence for a bottom-up mechanism of the theta-gamma code in the visual cortical networks (31, 32). Speculatively, the prefrontal and medio-temporal system, in concert with the oculomotor and the visual system may implement a mnemonic sampling loop. Novel perceptual information, reflected in gamma bursts, may be sampled bit by bit, at a theta pace (6), in order to integrate them into existing neuronal networks and to establish a continuous update of internal representations of the external world (2). Specifically, theta-gamma PAC may provide an optimal temporal code for long-term potentiation processes in the medial temporal lobe (20), the core system for episodic memories (21, 22), a coherent representation of time and space (33).

To conclude, this is the first study to demonstrate that a rhythmic visual brain stimulation can entrain complex cognitive functions, such as memory encoding, in the wake human brain. Specifically, memory entrainment was explained by participants responsiveness to a visual stimulation at a theta pace and the present findings underline a functional role of theta-gamma coupling as key mechanism in memory formation. More generally, the present findings show that sensory stimulation techniques open up new avenues for the investigation and the selective enhancement of functional mechanisms that underpin human cognition.

## Materials and Methods

### Subjects

20 university students (19 females, *M_age_* = 21.7, *SD_age_* = 3.6) voluntarily participated and received monetary reward or course credits. None of the subjects reported any history of neurologic or psychiatric disorders and all subjects had normal or corrected to normal visual acuity. Informed consent was obtained from all subjects and the experimental procedure agreed with the World Medical Association’s Declaration of Helsinki. All participants were included in the analyses.

### Stimuli and Procedure

#### Individual frequency adjustment

We conducted a pre-study to identify individual theta and alpha frequencies, for procedural details see (12). In the pre-study subjects saw 300 colored object pictures and made living/nonliving judgements. Individual frequencies were then selected based on the grand mean of each participant (all trials of all conditions). Mean frequencies for 20 subjects that returned to the lab for the main study were 5.4 Hz (SD = 1.1 Hz) for theta and 10.2 Hz (SD = 1.7 Hz) for alpha. We also used the pre-study to determine the gamma range, peaking at 50 – 80 Hz over posterior electrodes.

#### Stimuli

The stimuli were 600 colored object pictures (e.g. plants, animals, clothes, tools), taken from a standard library (Hemera Photo Objects). Pictures were presented at the center of a 19 in. computer screen at a visual angle of about 6.2 × 6.2°.

#### Experimental paradigm

During encoding subjects saw 450 objects, presented in one of three conditions: individual theta, individual alpha, or static, presented in a random order. In the theta and alpha conditions, the object was presented at the individual frequencies determined in the pre-study (see above) by controlling the presentation at every refresh cycle of a 72 Hz CRT monitor (one refresh cycle = 13.89 ms). For example, to establish a flicker rate of 6 Hz, the object was presented at a duty cycle of 6:6, i.e. six on and six off cycles. Each picture was presented following a black screen (1 s) and a white fixation dot (variable duration of 0.5 – 1 s), for 3 s, ending at a full duty cycle.

To maintain attention during the whole stimulus presentation, subjects had to detect a magenta-colored dot that appeared for 111 ms on 15% of the encoding trials at a random time and position and respond as quickly as possible by a button press. These stimuli were not presented during retrieval and excluded from further analyses.

The retrieval phase followed a 15 min filler task (solving simple math equations). 405 pictures from encoding were randomly intermixed with 150 new stimuli. Objects were presented for 2 s and subjects judged whether they would remember an object explicitly, with details from encoding (“remember”), the object seemed familiar (“know”), or the object was novel to them (“new”), i.e., remember/know paradigm; (29).

#### Behavioral analysis

Given the within-subject design, we compared the pure response rates of the three stimulation conditions: subsequently remembered (SR), subsequently known (SK), and subsequently forgotten (SF) responses. Note that the response rates do not need to be corrected by false alarm rates due to the within design. Our main comparison relied on the SR and SF responses and the subsequent memory effect (SME = SR-SF) for the theta and the alpha stimulation. Given the clear direction of the hypothesis that theta SSVEPs would lead to a higher SME than alpha SSVEPs, a one-sided test was applied for this main comparison. We quantified the overall memory entrainment effect (theta SME – alpha SME), to test if memory entrainment would be explained by the theta or the alpha stimulation. Theta and alpha SSVEPs were summarized by a mean z-score of the SSVEP signal at the peak electrode and the spread of the SSVEP signal across the scalp (across all 128 electrodes; Shannon entropy on SSVEP values, normalized from 0 to 1).

### Electroencephalogram (EEG) Apparatus and Analysis

#### Apparatus

EEG was recorded from 128 active electrodes using a BioSemi Active-Two amplifier system at 512 Hz, in a electromagnetically-shielded room. Two additional electrodes (CMS: Common Mode Sense and DRL: Driven Right Leg; cf. http://www.biosemi.com/faq/cms&drl.htm) served as reference and ground.

#### Preprocessing

EEG signals were band-pass filtered from 1 Hz to 120 Hz and segmented into epochs from -1500 ms to 4500 ms, regarding the stimulus onset. Eye-blinks and muscle artifacts were detected using an independent component procedure and removed after visual inspection. Noisy trials were identified visually and discarded (approx. 5 % - 10 %). We applied a correction of saccade-related transient potentials (34) to remove artifacts caused by miniature eye-movements (35), before signals were re-referenced to the average reference.

#### Steady state visually evoked potentials (SSVEPs)

The main analyses focused on the SSVEPs elicited by the theta and alpha stimulation. Specifically, the evoked spectral power was obtained by Morlet’s wavelets with approximately seven cycles at a resolution of 0.5 Hz for theta and alpha and a 2.0 Hz resolution in the gamma range. We identified the individual theta and alpha peak SSVEP signals, based on the grand mean spectral activity, and used the gamma range of 50-80 Hz, identified in the pre-study. Event-related changes for each condition and frequency were then calculated as the relative signal change of the post stimulus spectral activity, in percent, relative to a 500-250 ms for low frequencies, or 200-100 ms for gamma. For the analysis of theta and alpha SSVEPs as well as the gamma power in the SSVEP conditions, the SSVEP peak electrode of each condition was used (Figure 2).

#### Phase-amplitude coupling (PAC)

To assess PAC between the phase of both low frequencies, theta and alpha, and the gamma amplitude (illustrated in Figure 3B), the modulation index (MI) (16) was calculated as described previously (12). Specifically, the individual theta and alpha SSVEPs at the occipital peak electrode were filtered +/− 0.5 Hz around the individual peak frequency, and the 50-80 Hz filtered gamma signal, at all 128 electrodes. The MI was then calculated between the theta phase of the peak electrode in combination with the gamma amplitude, separately for each electrode, for the 500-3000ms time window. With the same procedure and using the same electrodes, we tested theta-gamma PAC across a 40-120Hz gamma range, in 5Hz steps, using a 20Hz sliding window for the gamma frequency, separated between conditions. The resulting MI value was averaged across all electrodes. For a circular plot of the subsequent memory differences (SR-SF) for gamma amplitudes, we z-standardized the gamma amplitudes across the 18 bins for each participant and condition. The SME difference (SR-SF) in standardized gamma amplitudes was averaged over the left temporal peak electrodes of the MI difference plot (Figure 2B).

## Acknowledgements

We thank Nikolai Axmacher and Malte Wöstmann for providing comments on an earlier version of the manuscript.

